# Temporal asymmetries and interactions between dorsal and ventral visual pathways during object recognition

**DOI:** 10.1101/2022.09.17.508376

**Authors:** Vladislav Ayzenberg, Claire Simmons, Marlene Behrmann

## Abstract

Despite their anatomical and functional distinctions, there is growing evidence that the dorsal and ventral visual pathways interact to support object recognition. However, the exact nature of these interactions remains poorly understood. Is the presence of identity-relevant object information in the dorsal pathway simply a byproduct of ventral input? Or, might the dorsal pathway be a source of input to the ventral pathway for object recognition? In the current study, we used high-density EEG – a technique with high temporal precision and spatial resolution sufficient to distinguish parietal and temporal lobes – to characterize the dynamics of dorsal and ventral pathways during object viewing. Using multivariate analyses, we found that category decoding in the dorsal pathway preceded that in the ventral pathway. Importantly, the dorsal pathway predicted the multivariate responses of the ventral pathway in a time-dependent manner, rather than the other way around. Together, these findings suggest that the dorsal pathway is a critical source of input to the ventral pathway for object recognition.

## Introduction

The primate visual system has historically been segregated into at least two independent visual pathways. The ventral visual pathway, which projects along the inferior temporal cortex (IT) and is involved primarily in visual object recognition, and the dorsal visual pathway, which projects along the parietal cortex and is involved primarily in visuospatial processing in the service of visually guided action. Despite their anatomical and functional differences, accumulating evidence suggests that the dorsal pathway may contribute to object recognition processes through its interactions with the ventral pathway. Yet, few studies have examined the precise temporal dynamics of object processing in dorsal and ventral pathways, leaving the directionality of these interactions unclear.

The dorsal pathway exhibits many functional properties that are crucial for object recognition. For instance, fMRI studies have shown that, like the ventral pathway, the dorsal pathway exhibited greater univariate activation to intact object images compared to scrambled object images, even when the objects did not afford action (Freud, Culham, et al., 2017; Grill-Spector et al., 1998). Moreover, the multivariate responses of dorsal regions were sufficient to decode an object’s category across variations in the object’s image-level properties and across variations among a category’s exemplars (Bracci & Op de Beeck, 2016; Vaziri-Pashkam et al., 2019). Indeed, object classification accuracy in dorsal regions matched, and sometimes surpassed, classification accuracy of ventral object regions (Ayzenberg & Behrmann, 2022b; Jeong & Xu, 2016). Other studies have shown that applying transcranial magnetic stimulation (TMS) over the dorsal pathway impaired aspects of object perception, such as configural processing and global shape perception (Romei et al., 2011; Zachariou et al., 2017; Zaretskaya et al., 2013). And, indeed, many patient studies have shown that damage to the dorsal pathway can impair perception of global shape (Dalrymple et al., 2007; Karnath et al., 2000; Riddoch et al., 2008; Thomas et al., 2012). Thus, the dorsal pathway not only represents information crucial for action, but also represents identity-relevant object properties crucial for recognition (Ayzenberg & Behrmann, 2022a; Freud et al., 2016).

There is also a high degree of anatomical and functional connectivity between dorsal and ventral pathways (Baizer et al., 1993; Kravitz et al., 2013; Takemura et al., 2016; Webster et al., 1994), suggesting that these pathways interact. However, the critical question regards the directionality of these interactions. Some studies have proposed that the ventral pathway transmits object information to the dorsal pathway to specify the action affordances of objects (Almeida et al., 2013; Garcea & Mahon, 2014; Xu, 2018). Indeed, regions of the left parietal cortex (in right-handed participants) exhibited preferential functional connectivity with the ventral pathway when participants viewed images of tools – a category with a high-degree of affordance information (Chen et al., 2017; Garcea & Mahon, 2012). On this view, the dorsal pathway does not contribute to object recognition *per se* but represents identity-relevant object properties as a byproduct of ventral input.

An alternative proposal, however, is that the dorsal pathway computes identity-relevant properties independently of the ventral pathway and may even propagate object information to the ventral pathway to support recognition (Ayzenberg & Behrmann, 2022a). Consistent with this proposal, electrophysiological recordings from both patients and monkeys have revealed that object information can be decoded in the dorsal pathway prior to that in the ventral pathway (Janssen et al., 2008; Regev et al., 2018). Moreover, temporary inactivation of the posterior parietal cortex (PPC) in monkeys reduces activation in ventral regions during object perception tasks (Van Dromme et al., 2016). Other research has shown that patients with extensive ventral damage nevertheless showed preserved responses to object shape in PPC, as well as an ability to recognize objects in certain contexts (Freud, Ganel, et al., 2017; Holler et al., 2019; Riddoch & Humphreys, 1987; Riddoch et al., 2008), suggesting that dorsal representations of shape are independent of the ventral pathway.

In a recent paper, Ayzenberg and Behrmann (2022b) used fMRI to characterize the functional contributions of the dorsal pathway to the ventral pathway during object recognition. They found that object decoding performance in shape-selective regions of PPC was comparable to decoding performance in the lateral occipital cortex (LO) – a ventral region crucial for object recognition (Behrmann et al., 2016; Grill-Spector et al., 2001; Pitcher et al., 2009). Additionally, they found that the multivariate response of the dorsal pathway mediated representations of shape in the ventral pathway, with effective connectivity analyses revealing that PPC signals predict the time course of LO activity, rather than the other way around. However, because fMRI has low temporal resolution, it is not possible to draw strong conclusions about the directionality of these interactions.

In the current study, we measure the temporal dynamics of the dorsal and ventral pathways using high-density electroencephalography (HD-EEG) – a neuroimaging method with millisecond precision and sufficient spatial resolution to localize dorsal and ventral pathways. Specifically, we selected channels of interest anatomically, corresponding to PPC and LO, and examined the directionality of their interactions. Although a limitation of EEG is its low spatial resolution, many studies have shown that the spatial resolution of HD-EEG is sufficient to dissociate lobes of the brain (Ferree et al., 2001; Hedrich et al., 2017; Koessler et al., 2009). Indeed, concurrent fMRI and EEG recording have shown that HD-EEG can be used to localize retinotopic responses in visual cortex to within 8 mm of the fMRI responses (Im et al., 2007). Moreover, because PPC and LO are situated on the surface of the brain, recordings from these regions are less impacted by noise, or other perturbations associated with distance, than deeper regions (Krishnaswamy et al., 2017). Thus, although the spatial resolution of HD-EEG is coarse when compared with other neuroimaging methods, it is sufficient to dissociate between neural activity in dorsal and ventral pathways.

We used an existing HD-EEG dataset in which participants were shown four categories of objects: tools, non-tools, insects, and birds (see Figure 1; Gurariy et al., 2022). In their initial study, Gurariy et al. (2022) used multivariate analyses and source localization to demonstrate comparable decoding of object shape and object category information in both dorsal and ventral pathways. However, they did not examine the temporal interactions between the pathways. Here, we characterized the object classification accuracy of each region across time and tested whether object category information could be decoded earlier in dorsal or in ventral areas. Moreover, we used Granger causality and time generalization analyses to examine whether the dorsal pathway predicted the time course of responses in the ventral pathway, or vice versa. To foreshadow our results, we found that category decoding in the dorsal pathway preceded the ventral pathway, and that the dorsal pathway predicted the response of the ventral pathway in a time-dependent manner. Together, these findings suggest that the dorsal pathway is a critical source of input to the ventral pathway for object recognition.

**Figure 1.**
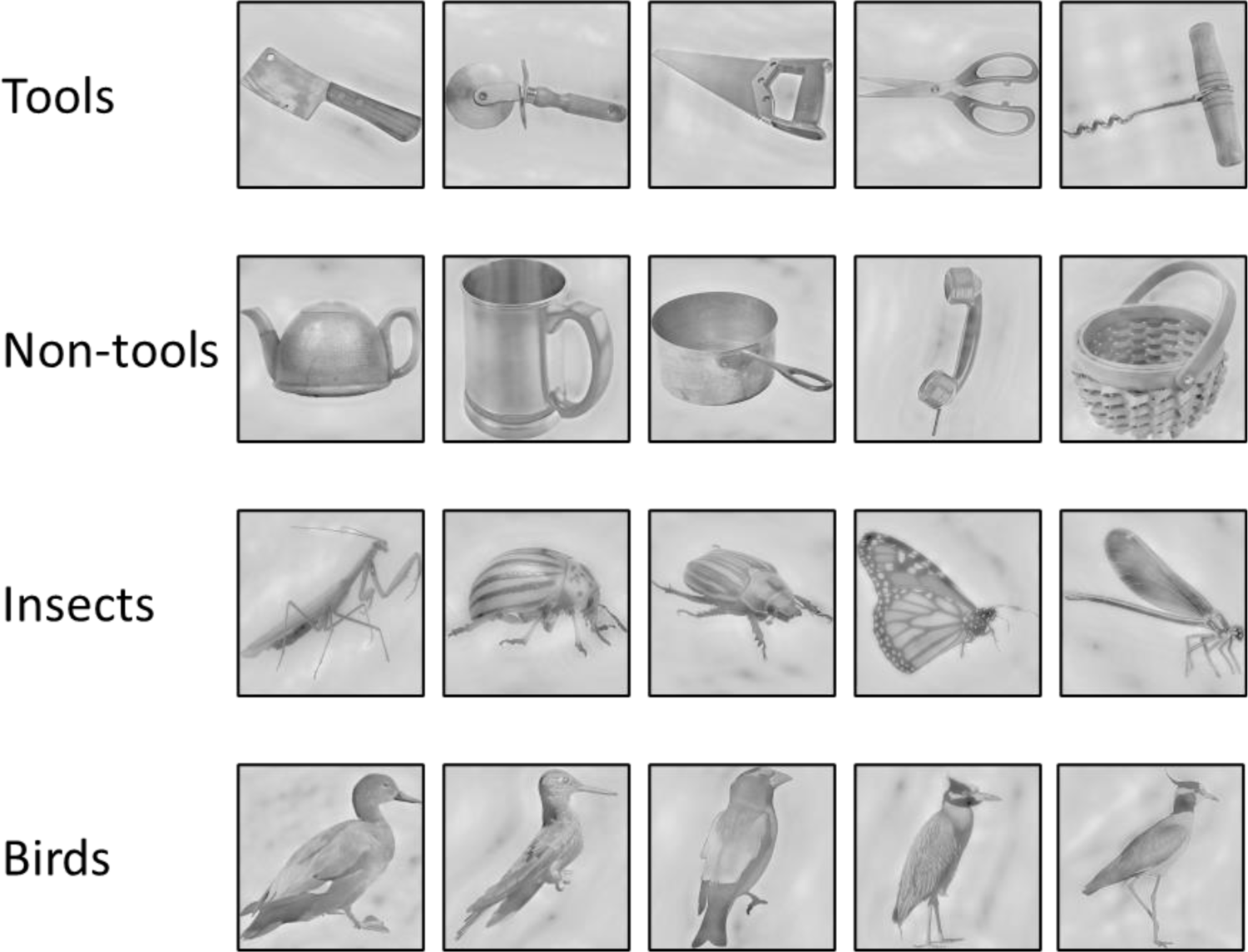
Stimuli presented to participants. Stimuli were comprised of four categories: tools, non-tools, insects, and birds. See Gurariy et al. (2022) for full details about stimulus design and presentation.

## Methods

### Participants

For a full description of the methods and procedures, see Gurariy et al. (2022). Twenty right-handed neurotypical adults (12 men, ages 18–38 years) with normal or corrected-to-normal visual acuity participated in the study. Each participant provided informed written consent. All protocols received approval by the Institutional Review Board at the University of Nevada, Reno. We received fully deidentified data from the original investigators.

### Apparatus

Stimuli were displayed on a Mitsubishi Diamond Pro270 CRT monitor (20 in., 1024 × 768) with a 120-Hz refresh rate, running via a 2.6-Mhz Mac Mini and presented using the PsychToolbox (Kleiner et al., 2007; Brainard, 1997; Pelli, 1997) for MATLAB (MathWorks Inc., 2007). Participants were seated 57 cm from the screen.

### EEG Data Acquisition

The EEG signal was continuously recorded using a 256-channel HydroCel Geodesic Sensor Net via an EGI Net Amps Bio 300 amplifier (Electrical Geodesics Inc.) sampling at 1000 Hz. The digital data were recorded using Netstation 5.0(1) software. Impedance values were kept at or below 100Ω. A photodiode was used to validate frame-accurate timing of stimulus presentation.

### EEG Preprocessing

EEG data were preprocessed using the EEG Lab Toolbox (Delorme & Makeig, 2004) and custom scripts written in MATLAB. Data were referenced from Cz to the algebraic average of all electrodes. A high bandpass of 0.1 Hz was applied to remove slow drift and electrical noise. Data were then downsampled from 1000 Hz to 250 Hz. Next, a low bandpass filter of 40 Hz was applied. Bad EEG channels were rejected using the ‘pop_rejchan’ and ‘pop_clean_rawdata’ functions in EEG Lab by identifying channels with extended flat line recordings and minimal channel correlation using joint probability of the recorded electrode. Independent component analysis (ICA) was used to identify and remove residual blink artifacts. Across participants, an average of 15.2 % of channels were removed.

The filtered time series was then segmented into 550-msec epochs (a 50-msec pre-stimulus baseline followed by 500 msec of electrophysiological data after stimulus onset). Event segmentation was performed using trigger markers that were sent to the acquisition computer at the onset of each trial. The temporal offset that existed between the physical presentation of the stimulus and the registration of the stimulus marker in the acquisition computer was measured using a photodiode and corrected for during trial segmentation. Following event segmentation, artifacts were detected using EEG Lab functions (pop_eegthresh) which rejects artifacts using thresholding. The EEG epochs described above for each participant were grouped into four conditions: tools, non-tools, insects, and birds.

### Channel Selection

Anatomically defined channels of interest were selected a priori in regions corresponding to PPC and LO using well-established cortical projections of EEG sensors (Koessler et al., 2009; Luu & Ferree, 2005). Specifically, channels were identified by first overlaying PPC (IPS0 and IPS1; Wang et al., 2014) and LO (Julian et al., 2012) probabilistic masks on a standard MNI brain with the Juliech Atlas, and determining the corresponding Brodmann areas for each (see Figure 2A). Channels corresponding to PPC (BA 7 and superior 19) and LO (BA 18 and inferior 19) were then identified using a template brain with electrodes arranged in the 10-10 system (Koessler et al., 2009). PPC electrodes consisted of 14 channels in the left hemisphere located approximately between P3 and Pz locations and 14 channels in the right hemisphere located approximately between Pz and P4 locations (see Figure 2B). LO electrodes consisted of 14 channels in the left hemisphere located approximately between the T5 and O1 locations, and 14 channels in the right hemisphere located approximately between T6 and O2 locations (see Figure 2B). For comparison, we additionally selected electrodes corresponding to occipital cortex, which consisted of 14 channels total approximately located between O1 and O2 locations. We also selected electrodes corresponding to frontal cortex, which consisted of 14 channels in the left hemisphere located approximately between F7 and Fz and 14 channels in the right hemisphere located approximately between Fz and F8 locations.

**Figure 2.**
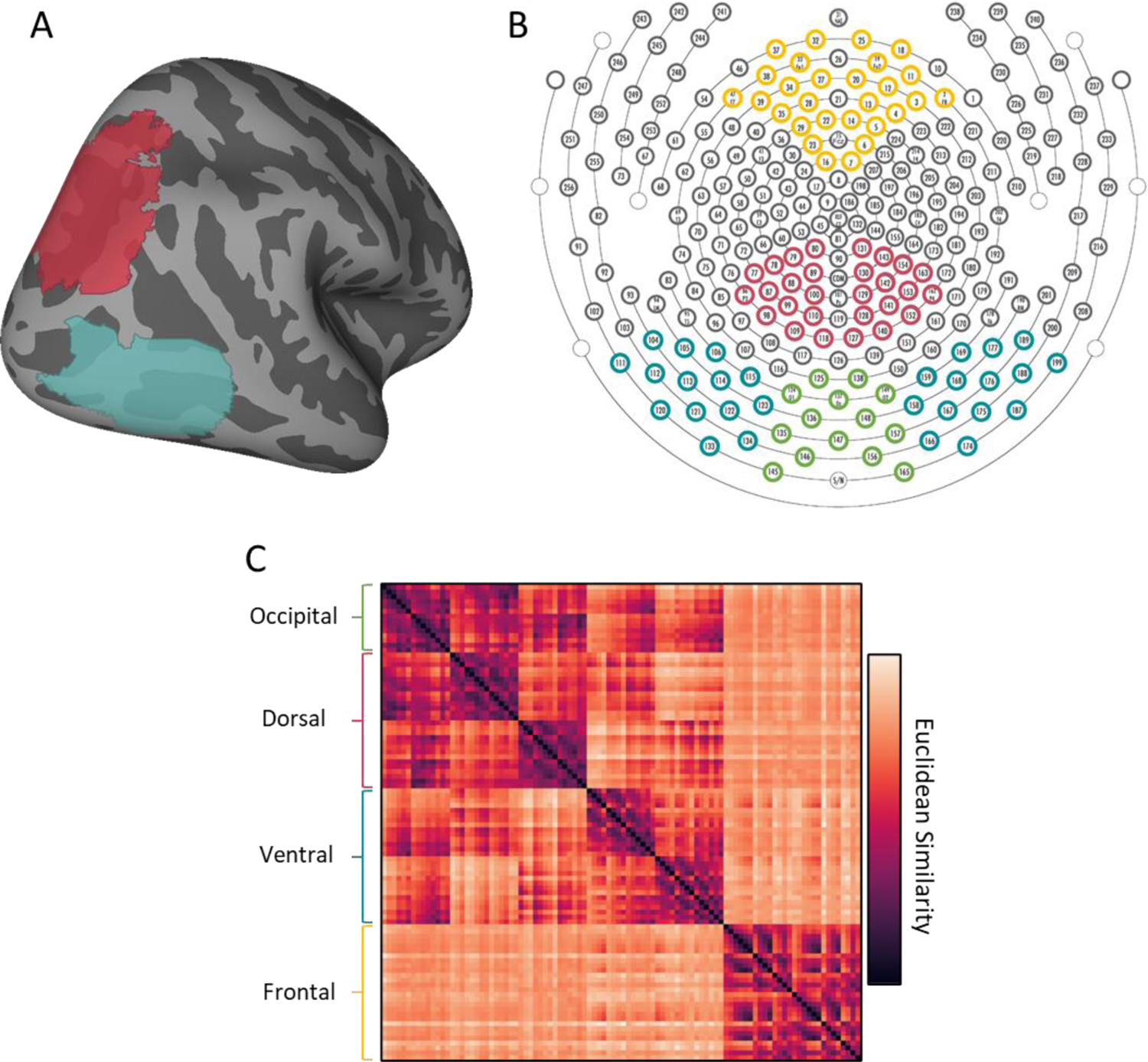
Anatomical regions of interest and their corresponding HD-EEG channels. (A) Probabilistic parcels for (red) PPC and (blue) LO projected onto an inflated standard brain (right hemisphere). (B) A diagram illustrating the channels of interest corresponding anatomically to (red) PPC and (blue) LO, as well as (green) occipital and (yellow) frontal control region. (C) A cluster map illustrating the similarity between channels-of-interest. Hierarchical clustering of EEG channels revealed that dorsal channels were functionally more similar to one another than to other channels-of-interest; likewise, ventral channels were functionally more similar to one another than other channels-of-interest.

To further validate our channel selection decisions, we tested whether our anatomically defined occipital, dorsal, ventral, and frontal channels are functionally distinct. Given the diffuse nature of the EEG signal, the goal of this analysis is to test whether the functional profile of each region was unduly affected by signals from elsewhere in the brain. Principal component analysis (PCA) was conducted on all channels for each participant, and we retained components that explained 95 % of the variance for each participant – resulting in a unique number of components for each participant. Then, we tested whether occipital, dorsal, ventral, and frontal channels load onto distinct components. Specifically, we calculated the Euclidean distance between the component loadings for each channel with every other channel in a pairwise fashion, providing a metric of functional similarity between all channels. Hierarchical cluster analyses of the mean participant similarity matrix revealed that our anatomically selected occipital, dorsal, ventral, and frontal channels are more similar to one another than to the other regions (Figure 2C). Thus, as consistent with previous work (Koessler et al., 2009), HD-EEG provides sufficient spatial resolution to decode signals from distinct lobes of the brain with minimal impact from other neural sources.

Finally, to ensure that any differences between regions could not be accounted for by different levels of noise, we also computed the signal-to-noise ratio (SNR) for each region. SNR was computed using two methods. We first computed overall SNR for the entire time course by dividing the mean signal by the standard deviation of the time course. This analysis method revealed no differences in SNR between regions (*ps* > .090; occipital: 6.26, dorsal: 4.85, ventral: 5.59, frontal: 6.19). In the second approach, we computed SNR by dividing the mean signal during the stimulus period by the standard deviation of the pre-stimulus baseline period. As was true of the first approach, this analysis revealed no significant differences between regions (*ps* > .335; occipital: 0.65, dorsal: 0.57, ventral: 0.65, frontal: 0.70). Thus, the equivalence of the SNR between regions indicates that any differences we obtain are not a function of signal strength in any lobe or of differential noise.

### Stimuli and procedure

The stimuli comprised four categories, each consisting of five exemplars (resulting in 20 unique images; see Figure 1). The categories included two inanimate classes: tools, here defined as objects that elicit a motor program specific to the object’s function (e.g., twisting for the corkscrew), as well as non-tool objects, here defined as objects that are manipulable, but do not elicit a motor program specific to the object’s function (e.g., all non-tools elicit grasping) (Garcea & Mahon, 2012). There were also two animate classes, birds and insects. All stimuli were processed using the SHINE toolbox (Willenbockel et al., 2010) to control for low-level differences in luminance and spatial frequency.

On each trial, an image was presented at the center of the screen (15°× 15°) for 300 msec followed by an interstimulus interval (ISI) lasting between 800 and 1200 msec. Each of the 20 exemplars that made up the four categories was presented 84 times resulting in 420 trials per category and 1680 trials in total. The order of presentation was randomized. To maintain their attention, participants performed an orthogonal task wherein they pressed the space bar if an image appeared at a reduced luminance (50% reduction), and this occurred on 5% of the trials. Reduced luminance trials were excluded from all subsequent analysis.

### Data analysis

#### Category decoding

For each participant, region and timepoint, a support vector machine (SVM) classifier was trained on the multivariate pattern for three exemplars from each category, and then required to predict the category of the two left out exemplars (100-fold cross-validation). Above chance (0.25) decoding accuracy for each timepoint was determined using a bootstrap procedure, wherein participants’ decoding data was resampled 10,000 times (with replacement) and 95% bootstrap confidence intervals were constructed around the classification accuracy for each timepoint. Qualitatively similar results were found when using different classifiers (e.g., Naïve bayes) or statistical tests (e.g., one-sample t-test comparisons to chance).

#### Effective connectivity

For each participant and region, the channel × time (stimulus onset to offset) response for each object was concatenated with one another to form one continuous timeseries. A single null value was inserted between every object’s timeseries to prevent prediction of temporally discontinuous timepoints. Multivariate Granger causality analyses (Barnett & Seth, 2014; Barrett et al., 2010) were then conducted twice, once with the dorsal pathway channels as the predictor and once with the ventral pathway channels as the predictor. Following prior work (Roebroeck et al., 2005), effective connectivity between the areas was calculated by subtracting the dorsal → ventral *F* statistic from the ventral *→*dorsal *F* statistic. Multivariate Granger causality was tested with a maximum lag of 50 ms, with the best fitting lag for each direction used to compare dorsal and ventral pathways. Following previous work, a group analysis was conducted using a Wilcoxon signed-rank test comparing *F*-difference values to 0.

#### Time generalized representational similarity analyses

For each participant and region, a 20 × 20 representational dissimilarity matrix (RDM) was created for every time point by computing the Cosine similarity between the multivariate channel responses for one object with every other object in a pairwise fashion. RDMs were then averaged across participants separately for PPC and LO. Then, for every timepoint, the upper triangle of the resulting matrix (including the diagonal) for the dorsal pathway was correlated with the RDM of every timepoint of ventral pathway. To test whether the dorsal pathway predicted the ventral pathway, or the other way around, we subtracted the correlations below the diagonal (dorsal → ventral) from the correlations above the diagonal (ventral → dorsal). Specifically, for each participant, we subtracted each timepoint from the timepoint with the equivalent coordinates mirrored along the diagonal. For example, the timepoint with coordinates 100 ms in the dorsal pathway (*y*-axis) and 120 ms in the ventral pathway (*x*-axis) was subtracted from the time point with coordinates 120 ms in the ventral pathway and 100 ms in the dorsal pathway. A significant positive difference from 0 would indicate that the dorsal pathway better predicts future times points of the ventral pathway, rather than the other way around. The mean difference across all correlation subtractions was taken for each participant, and submitted to a one-sample t-test comparison to chance (0).

## Results

### The time course of object classification

We first examined the time course of identity-relevant object information in dorsal and ventral pathways. Specifically, we tested when the multivariate pattern in each area could be used to decode the objects’ categories (see Figure 3). We found that peak decoding accuracy was comparable in both pathways, with the peak accuracy at 210 ms in the dorsal pathway (35.2%) and 246 ms in ventral pathways (34.9 %). But the critical question has to do with the temporal signature, namely, in which cortical area was object information decoded earlier in time? An analysis of the first onset of significant decoding found that object category information could be decoded above chance earlier from the dorsal pathway (28.5 %) at 66 ms, whereas category information was decoded from the ventral pathway (29.1%) at 94 ms, a difference of 28 ms (see Figure 3A).

**Figure 3.**
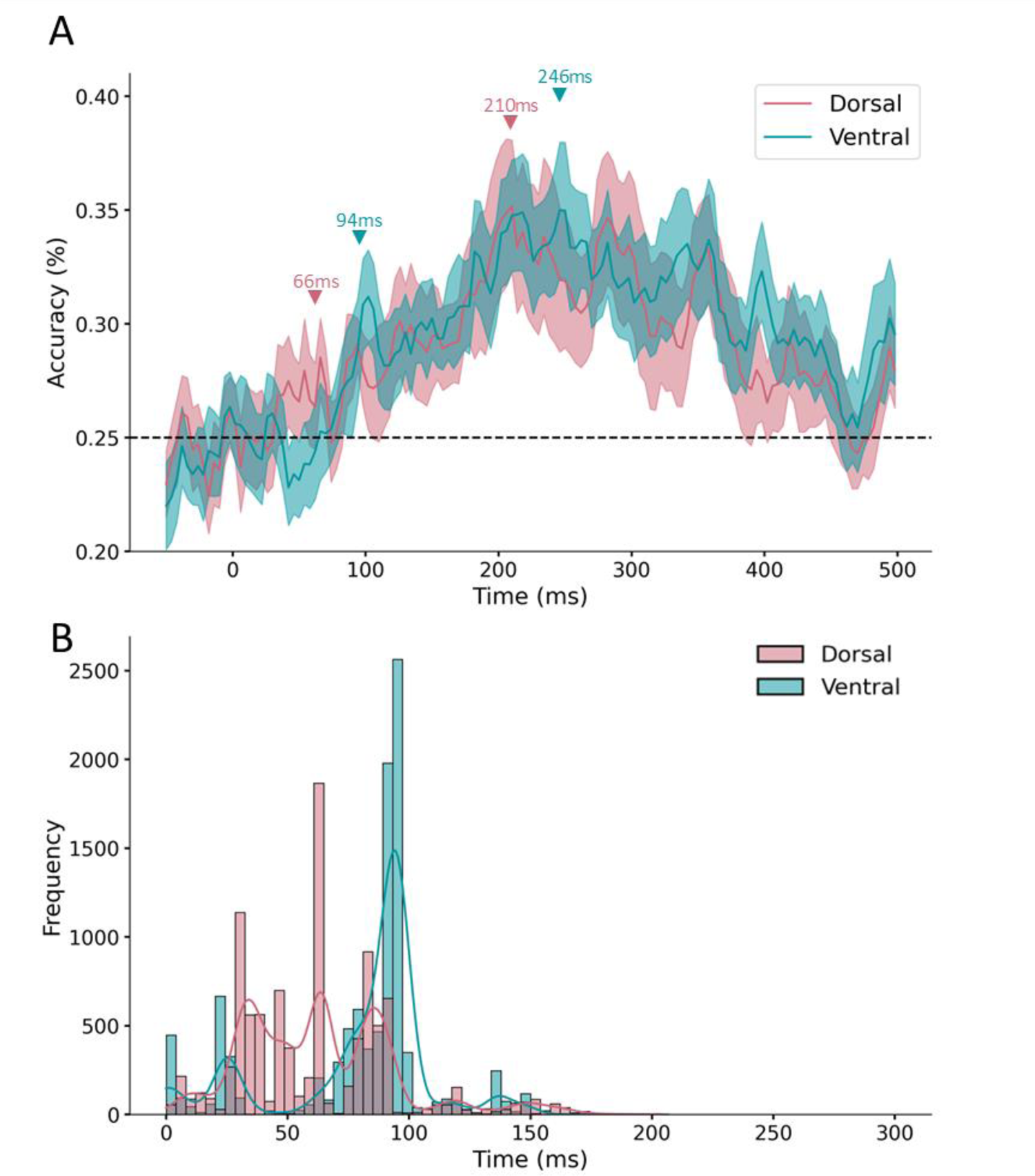
Results of the decoding analyses. (A) Time course of category decoding for (red) dorsal and (blue) ventral channels. Classification accuracy is plotted along the y-axis as a function of time (ms). Data are plotted from the pre-stimulus period to post-stimulus period (−50 ms to 500 ms). Shaded regions indicate standard error of the mean (SE). (B) A histogram illustrating the earliest onset of above chance classification for 10,000 resamples for (red) dorsal and (blue) ventral channels. The y-axis illustrates proportion with which each time point exhibited above chance classification accuracy. Data are plotted for the stimulus period (0 ms to 300 ms).

We next tested whether these timing differences between dorsal and ventral pathways were statistically consistent. Participant data from dorsal and ventral pathways was resampled 10,000 times (with replacement). On each resample a one-sided, one-sample t-test was conducted for every timepoint of participants’ data, and the first timepoint with above chance decoding was recorded. Again, to minimize spuriously significant timepoints, only those timepoints with significant decoding for at least two consecutive timepoints were recorded (results were qualitatively similar without this step).

Across resamples of the data, the decoding in the dorsal pathway occurred at a median time of 66 ms, whereas decoding in the ventral pathway occurred at a median time of 94 ms, a difference of 28 ms. A Binomial comparison to chance (0.50) revealed that the dorsal pathway preceded the ventral pathway in 70.1 % of resamples, significantly more often than would be predicted by chance (0.50; *p* < 0.001; Binomial test; see Figure 3B). Altogether, these results show that identity-relevant object information is present in the dorsal pathway prior to the ventral pathway.

For comparison, we also examined the presence of identity-relevant object information in occipital cortex and frontal cortex. Occipital cortex was chosen because it is considered the earliest stage of visual processing in the cortex and has direct connections to dorsal and ventral visual pathways. Hypothetically, then, identity-relevant information in the dorsal pathway may reflect propagation from occipital cortex, rather than a computation of the dorsal pathway per se. Frontal cortex was also examined because it has been suggested to be a source of feedback to the ventral pathway (Bar et al., 2006; Kar & DiCarlo, 2021) and, hypothetically could even be the source of identity information in the dorsal pathway.

These analyses revealed above chance decoding at 66 ms in occipital cortex (same time as dorsal) and 110 ms in frontal cortex (after dorsal and ventral pathways). Resamples of the data revealed a median decoding time of 70 ms in occipital cortex and 106 ms in frontal cortex. Binomial comparisons to the dorsal pathway of these distributions revealed that decoding in the dorsal pathway preceded both occipital cortex (58.1 % of resamples; *p* < .001) and frontal cortex (77.7 % of resamples; *p* < .001). By contrast, decoding in the ventral pathway was preceded by occipital cortex (58.1 % of resamples; *p* < .001), and the ventral pathway preceded frontal cortex (72.3% of resamples; *p* < .001).

#### Control analyses

We found that category decoding in the dorsal pathway preceded the ventral pathway. However, given the relatively small differences in timing between areas (28 ms), it is possible that these differences are spurious and reflect the idiosyncrasies of the stimulus categories, the channels selected, or the influence of signals outside our regions of interest. We conducted several control analyses to address these concerns and investigate the consistency of identity-relevant information in dorsal and ventral pathways.

We first tested whether differences between dorsal and ventral pathways were driven by the inclusion of the tool category. It is well known that the dorsal pathway is particularly sensitive to objects that afford action, such as tools (Almeida et al., 2010; Chao & Martin, 2000) and, indeed, prior HD-EEG work has shown better decoding for tools than other categories in the dorsal pathway (Gurariy et al., 2022). Thus, it is possible that the differences in timing we observe are driven solely by the inclusion of tools. To test this possibility, we repeated our decoding analysis in the absence of the tool category (non-tools vs. birds vs. insects; chance = 0.33). Consistent with the results above, this analysis revealed that decoding again occurred earlier in the dorsal pathway (38.5%) at 50 ms than the ventral pathway (36.3%) at 98 ms, a difference of 48 ms.

We next tested whether any of the other categories could explain the timing differences between dorsal and ventral pathways. To this end, we conducted decoding analyses on dorsal and ventral pathways in each participant after iteratively removing each category one a time (e.g., decoding with birds or insects or tools or non-tools removed; chance = 0.33). To provide an overall measure of decoding consistency across all object iterations we resampled these data 1,000 (with replacement) and calculated the onset of first decoding on every resample. Across the aggregate of all resamples of the data (i.e., every iteration of stimulus categories for participants), we found that median category decoding in the dorsal pathway (58 ms) preceded the ventral pathway (86 ms) by 28 ms. A Binomial comparison of the distributions (0.50) revealed that this difference was statistically significant (*p* < .001), with dorsal preceding ventral in 66.2% of resamples. Together, these analyses suggest that our results are not driven by tool category specifically, nor variations in which stimulus categories are included in the decoding.

The next possibility we tested is whether differences between dorsal and ventral pathways are driven by the specific channels-of-interest we selected for each region. If this possibility is true, then slight differences to the channel selection criteria may alter the results. To test this possibility, we randomly resampled (with replacement) the channels-of-interest for each participant within each region 1,000 times and repeated our primary decoding analysis on every resample. This analysis revealed that, across resamples of the channels decoding in the dorsal pathway preceded the ventral pathway (medians = 66 ms vs. 98 ms), by a difference of 32 ms. A Binomial comparison of the distributions (0.50) revealed that this difference was statistically significant (*p* < 0.001), with dorsal preceding ventral in 89.3% of resamples. Thus, our decoding results are consistent across variations in the specific channels selected.

Finally, we examined whether our decoding results could be explained by signals outside dorsal and ventral pathways. Although our hierarchical cluster analyses showed that dorsal and ventral channels were functionally distinct from other regions (see Methods), the signals we measured over dorsal and ventral pathways are likely influenced by channels elsewhere in the brain. To test this possibility, we repeated our decoding analyses after controlling for the response of all channels outside of our two key regions of interest.

To this end, we first conducted PCA on the timeseries response for all the channels of the brain for every participant, while excluding dorsal and ventral pathways, and retained components that explain 95% of the variance. These components provide a multivariate summary of the functional responses of all channels outside of the dorsal and ventral electrodes. Next, for each participant, we regressed each channel corresponding to the dorsal and ventral pathway by the principal components from the rest of the brain. Finally, we repeated our decoding analyses using the residual responses of the dorsal and ventral channels. This analysis revealed significant decoding in the dorsal pathway at 50 ms, and significant decoding in the ventral pathway at 78 ms, a 28 ms difference. Thus, although our channels-of-interest were likely influenced by sources outside dorsal and ventral pathways, these external sources do not explain the differences in timing between dorsal and ventral pathways. Altogether, these findings suggest that our decoding results cannot be explained by variations in the stimulus categories, selected channels, nor sources outside our regions of interest. Instead, our findings are robust across all these variations.

### Effective connectivity analyses

The results above revealed that object classification in the dorsal pathway preceded that of the ventral pathway. However, it is unclear whether the dorsal pathway is also a source of input to the ventral pathway. Indeed, it is possible that object processing simply occurs in parallel in dorsal and ventral pathways with a temporal advantage for dorsal over ventral but with little, if any, interaction.

To test whether the dorsal pathway is a potential source of input to the ventral pathway, we conducted hypothesis-driven multivariate Granger causality analyses (Barnett & Seth, 2014; Barrett et al., 2010). The premise underlying Granger causality analyses is as follows. The dorsal pathway can be said to predict the response of ventral pathway if incorporating the past multivariate responses of dorsal pathway (i.e., *t*-1) improves the prediction of current multivariate responses of the ventral pathway over above ventral’s own past responses.

Granger causality analyses were conducted in both directions (dorsal → ventral and ventral → dorsal), with the resulting *F* statistics subtracted from one another. A Wilcoxon signed-rank one-sample *t*-tests comparisons to 0 revealed significant effective connectivity from the dorsal to the ventral pathway, *W*(19) = 160, *p* = .020, *d* = 0.52. This finding suggests that the multivariate time course of the dorsal pathway predicts the ventral pathway, rather than the other way around.

For comparison, we also examined the interactions between occipital and frontal cortices with dorsal and ventral pathways. These analyses revealed that occipital cortex was a significant predictor of the ventral pathway, *W*(19) = 181, *p* = .002, *d* = 0.72, but not the dorsal pathway (*p* = .378). We also found that the dorsal pathway was a significant predictor of frontal cortex, *W*(19) = 160, *p* = .005, *d* = 0.65, whereas the ventral pathway was not (*p* = .249).

### Time generalization analyses

If the dorsal pathway precedes and predicts future responses of the ventral pathway, then the strongest correlation between dorsal and ventral pathways should be between the present timepoint of the dorsal pathway with future timepoints of the ventral pathway. To test this prediction, we conducted time generalized representational similarity analyses (King & Dehaene, 2014; Kriegeskorte et al., 2008).

See Figure 4A for the time generalization matrix. Across participants, correlations were stronger when the dorsal pathway was correlated with future timepoints of the ventral pathway, than when the ventral pathway was correlated with future timepoints of the dorsal pathway, *t*(19) = 2.83, *p* = .011, *d* = 0.63.

**Figure 4.**
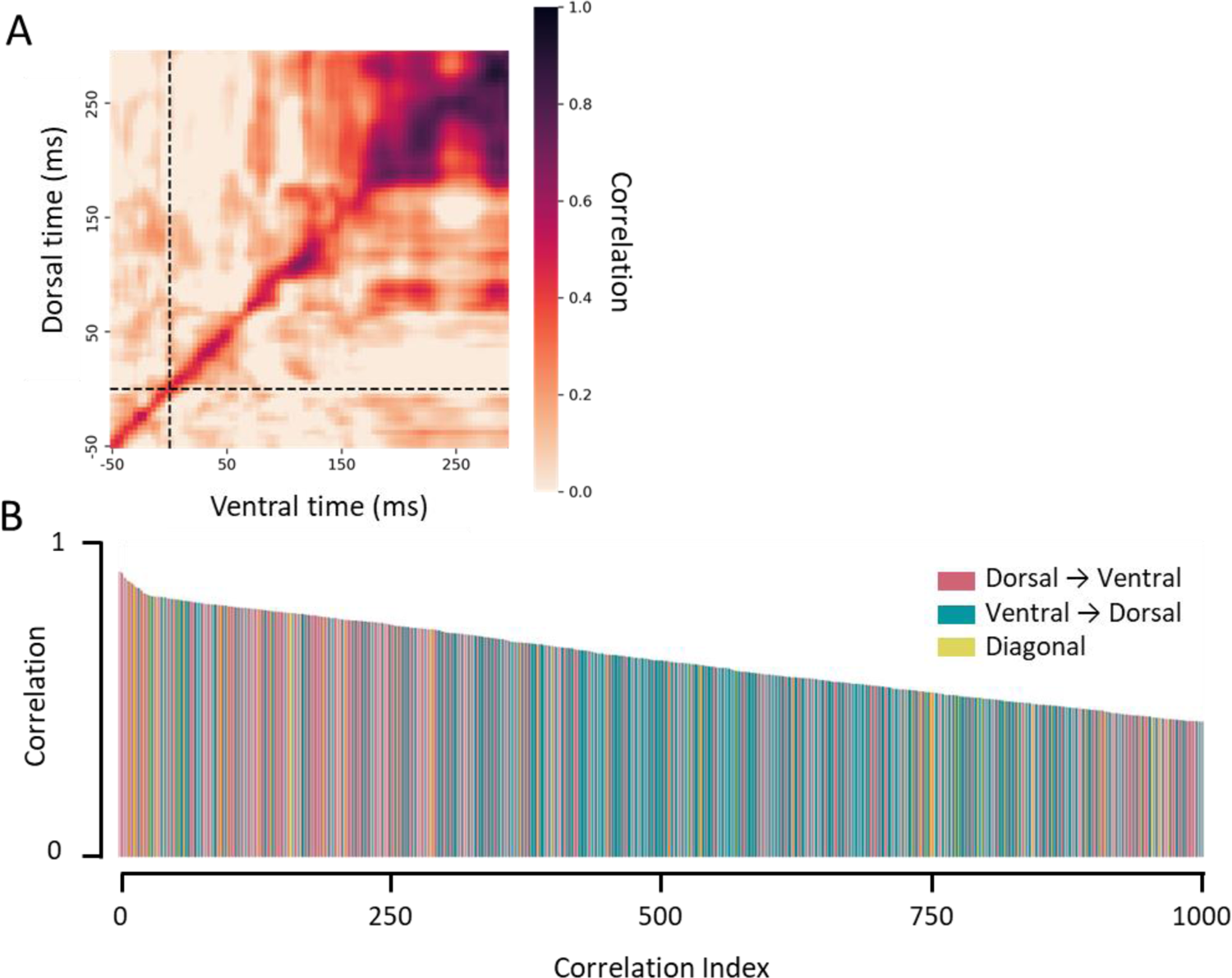
Time generalized representational similarity analyses. (A) Time generalization matrix illustrating the correlation between RDMs for each dorsal time point with each ventral time point. Correlations were stronger for dorsal → ventral correlations than ventral → dorsal correlations. Data are plotted from the pre-stimulus to stimulus offset period (−50 ms to 300 ms). Dotted line indicates stimulus onset. (B) Top 1000 correlations, rank ordered from highest to lowest for (red) dorsal → ventral correlations, (blue) ventral → dorsal correlations, and (yellow) correlations along the time diagonal.

To further visualize the patterns of interactions between dorsal and ventral pathways, we rank ordered the correlations from strongest to weakest and labeled each according to whether they reflected dorsal → ventral correlations, ventral → dorsal correlations, or correlations along the diagonal of the matrix (see Figure 4B). This visualization revealed that the strongest correlations were overwhelmingly those in which dorsal signals predicted ventral signals. Indeed, 9 of the 10 strongest correlations are those in which the dorsal pathway predicted ventral pathway, with the single remaining value reflecting a correlation along the diagonal of the matrix. The mean prediction delay between dorsal and ventral pathways was 8.8 ms.

To characterize this pattern quantitatively, we conducted a rolling Binomial analysis in increments of 50, where we tested whether dorsal → ventral correlations comprise a larger proportion of the correlations than would be predicted by chance (0.49). This analysis revealed that dorsal → ventral correlations comprised the majority of the top correlations up to the first 500 indexed correlations (55.2%, *p* = 0.006). At its peak, dorsal → ventral correlations comprised 66% of the top 300 correlations (*p* < 0.001), with a mean prediction delay of 34.3 ms. Altogether these results suggest that the dorsal pathway predicts the future response of the ventral pathway, rather than other way around.

Although these results suggest that the dorsal pathway propagates object information to the ventral pathway, it is unclear whether this effect reflects direct or indirect interactions. Indeed, as mentioned previously, other research has found that feedback to the ventral pathway may occur via frontal regions (Bar et al., 2006; Kar & DiCarlo, 2021). Thus, the dorsal pathway may interact with the ventral pathway by way of frontal cortex.

To test this possibility, we conducted partial correlation analyses controlling for the influence of frontal channels. As in the previous analysis, each timepoint of the dorsal pathway was correlated with each timepoint of the ventral pathway, however, here, the multivariate response of frontal cortex for each dorsal timepoint was included as a covariate. This analysis revealed that, across participants, partial correlations were strongest when the dorsal pathway was correlated with future timepoints of the ventral pathway, even when controlling for the influence of frontal cortex, *t*(19) = 2.85, *p* = .010, *d* = 0.64.

For completeness, we also examined whether the dorsal pathway predicted the ventral pathway, when, instead, we used the multivariate response of occipital cortex as a covariate. This analysis again revealed that the dorsal pathway was correlated with future timepoints of the ventral pathway, even when controlling for the influence of occipital cortex, *t*(19) = 4.14, *p* < .001, *d* = 0.93. Altogether, these findings are indicative of direct interactions between dorsal and ventral pathways, with dorsal propagating information to the ventral pathway, rather than the other way around.

Finally, we examined whether there are direct functional connections between occipital and frontal cortices with dorsal and ventral pathways. We found that occipital cortex predicted the response of the ventral pathway, *t*(19) = 3.03, *p* = .007, *d* = 0.68, but not the dorsal pathway (*p* = .78). We did not find any significant relations between frontal cortex with either dorsal or ventral pathways (*ps* > .226).

## General Discussion

In the current study, we examined the temporal dynamics and interactions of the dorsal and ventral visual pathways. Although an increasing number of studies have proposed that dorsal and ventral pathways interact during the process of object recognition, few studies have examined the timing and directionality of object processing in the two visual pathways. Using HD-EEG, we found that the dorsal pathway represented identity-relevant object information prior to the ventral pathway. Moreover, we found that the dorsal pathway predicted the response of the ventral pathway in a time-dependent manner. Altogether, these findings suggest that identity-relevant object information in the dorsal pathway arises independently of the ventral pathway and may be a source of input to the ventral pathway for object recognition.

However, a limitation of the current study is that EEG and the study design affords only coarse anatomical specificity. Although previous work has shown that HD-EEG can localize responses to within 8 mm of their anatomical location (Im et al., 2007; Koessler et al., 2009), the precise anatomical location of our channels of interest is not clear. Specifically, we used a standard brain template to select channels that encompass PPC and LO. Nevertheless, HD-EEG does have sufficient spatial resolution to dissociate lobes of the brain (Ferree et al., 2001; Hedrich et al., 2017; Koessler et al., 2009), allowing us to generally distinguish between dorsal and ventral pathways. Indeed, PCA of the EEG channels revealed that our anatomically-selected dorsal and ventral channels were functionally distinct from each other, as well as from other plausible neural sources (i.e., occipital and frontal cortex). Furthermore, our decoding and connectivity results remained consistent even when we controlled for signals from outside the dorsal and ventral pathway. Finally, it is important to note that the spatial limitations of HD-EEG should have attenuated any timing differences between dorsal and ventral pathways, not strengthened them. That we consistently found that the response of the dorsal pathway preceded and predicted the ventral pathway across a range of analyses suggests that our results are based on signals from distinct regions. Indeed, it is likely that our results underestimate the true temporal asymmetries and interactions between dorsal and ventral pathways. Nevertheless, future work using intracranial recording with patients (Regev et al., 2018), or concurrent scalp EEG and MRI recordings will be needed to strengthen these conclusions.

Although there are relatively few studies that compare the timing of object processing in dorsal and ventral pathways, our timing results are remarkably consistent with existing data. Specifically, in our primary analyses we found decoding in the dorsal pathway at 66 ms and in the ventral pathway at 94 ms. Similarly, electrocorticographic (ECoG) data from patients showed visual onsets in IPS at approximately 60 ms and at approximately 100 ms in LOC (Regev et al., 2018). Other work from monkeys show comparable timing in dorsal and ventral pathways, even when correcting for the differences in conduction time between monkey and human brains (Chen et al., 2006; Janssen et al., 2008). For instance, Janssen et al. (2008) found shape selective processing in the dorsal pathway of monkeys at 65 ms. Thus, although HD-EEG has relatively coarse spatial resolution, the timing of dorsal and ventral pathway in our study is consistent with studies using spatially precise techniques.

Why might category information arise earlier in the dorsal pathway? One possibility is that earlier onsets in the dorsal pathway reflect attentional processing. Attention is the process of selecting a target stimulus over competing distractors. It is well known that the dorsal pathway plays an important role in visual attention (Behrmann et al., 2004; Corbetta & Shulman, 2002), and thus the close proximity to cortical regions subserving attentional processing in the dorsal pathway may have provided the timing advantage we observed. Although additional experiments are needed to rule out this possibility definitively, on the basis of prior work and our own results, we believe this explanation is unlikely. First, classic electrophysiological studies in monkeys and EEG studies in humans typically find that the effects of visual attention occur after 100 ms which is much later in time than the decoding results we found here (Mehta et al., 2000; Posner et al., 1973; Schneider et al., 2012). Second, neurally, attention typically results in univariate increases in the response to attended stimuli (i.e., gain) and decreases to the unattended stimuli (McAdams & Maunsell, 1999; Wojciulik et al., 1998), as well as enhanced SNR for the attended stimulus (Mitchell et al., 2009). Given that SNR was matched in dorsal and ventral pathways in our data (and marginally higher in the ventral pathway; see Methods), it is not clear how an increased univariate response in the dorsal pathway would lead to earlier multivariate decoding of object category. In fact, univariate enhancement of identity-relevant properties in the dorsal pathway would require that these particular properties are already represented in the dorsal pathway. Relatedly, it is unclear how attention might account for our findings that the multivariate response of the dorsal pathway predicted the time-varying responses of the ventral pathway.

A more probable explanation for the observed dorsal and ventral timing results is that they reflect the differential input of magnocellular and parvocellular pathways to dorsal and ventral cortices, respectively (Almeida et al., 2013; Laycock et al., 2007). The magnocellular pathway is able to rapidly transmit coarse object information to the dorsal pathway via direct projections from subcortical regions, which partially bypass occipital cortex (Felleman & Van Essen, 1991); whereas the parvocellular pathway transmits higher resolution object information to the ventral pathway by way of occipital cortex (Bar et al., 2006; Collins et al., 2019; Wang et al., 2022). Consistent with these mechanisms, our results suggest that identity-relevant information in the dorsal pathway did not depend solely on input from occipital cortex. Specifically, we found that decoding in the dorsal pathway occurred as early or earlier than in occipital cortex, and that occipital cortex was not a significant predictor of the dorsal pathway. By contrast, decoding in occipital cortex preceded the ventral pathway and occipital cortex was a significant predictor of the time-varying responses of the ventral pathway. Importantly, TMS and fMRI studies suggest that the coarse information transmitted by the magnocellular pathway is sufficient to compute an object’s global shape structure (Wang et al., 2022), and that this information can support decoding of object category in the dorsal pathway (Ayzenberg & Behrmann, 2022b). Furthermore, studies have shown that such structural information may then be transmitted to the ventral pathway to support object recognition (Ayzenberg & Behrmann, 2022b; Bar et al., 2006; Kveraga, Boshyan, et al., 2007). Thus, the earlier decoding of object category in the dorsal pathway may be explained by privileged magnocellular input.

Although our results are most consistent with the dorsal pathway as a source of input to the ventral pathway, it is almost certainly the case that there are bidirectional interactions between the two pathways (Kravitz et al., 2011; Kravitz et al., 2013). For instance, at later stages of processing, the ventral pathway may provide semantic information about an object’s affordances to the dorsal pathway to help coordinate actions (Almeida et al., 2013; Chen et al., 2017; Garcea & Mahon, 2014). Moreover, identity-relevant object information may be transmitted in a recurrent manner between dorsal and ventral pathways as needed to accomplish either object recognition or coordinate motor actions. It is unlikely, however, that our results in the dorsal pathway reflect differential action affordances for each category. Specifically, decoding in the dorsal pathway preceded the ventral pathway even when we removed categories with a high degree of action affordance (i.e., tools). Moreover, the neural signals for action planning (including saccade execution) are typically found later in time than our decoding results here (e.g., > 100 ms; Cui & Andersen, 2011; Hamm et al., 2010; Handy et al., 2003). Instead, our findings suggest that the dorsal pathway provides an early source of input to the ventral pathway to support object recognition. Future work should examine how the directionality and dynamics of dorsal-ventral interactions change for different tasks and anatomical stages.

There is also evidence that dorsal and ventral pathways are situated within a broader network of regions that are involved in object recognition (Kravitz et al., 2013). For instance, accumulating evidence suggests that prefrontal cortex (PFC) is also a source of feedback to the ventral pathway during object recognition (Bar et al., 2006; Kar & DiCarlo, 2021). Specifically, coarse object information arises in PFC via the dorsal pathway (Kveraga, Boshyan, et al., 2007) and this information is then used to constrain the possible identity of the object (Kveraga, Ghuman, et al., 2007). Might dorsal and ventral pathways interact indirectly via frontal cortex? Our results suggest not. Specifically, we found that the dorsal pathway predicted the time-varying responses of the ventral pathway even when controlling for frontal channels – a finding consistent with research showing direct connectivity between dorsal and ventral pathways (Janssen et al., 2018; Takemura et al., 2016; Webster et al., 1994). Moreover, and contrary to prior work, we did not find significant functional connectivity between frontal cortex with either dorsal or ventral pathways. One explanation for this discrepancy is that input from frontal cortex is needed in challenging contexts when an object’s properties are difficult to see (Bar et al., 2006; Kar & DiCarlo, 2021). However, the stimuli in the current study were clearly visible and easy to identify.

The primary goal of our study was to explore the temporal dynamics and interactions of dorsal and ventral pathways. Yet, one might nevertheless wonder, what kind of information supports decoding of object category in the dorsal pathway? One possibility is that, in the current study, decoding was accomplished by relying on low-level visual properties. This possibility is unlikely because the stimuli were controlled for low-level properties like luminance and spatial frequency (see Methods). Another possibility is that very coarse object properties, such as elongation, were sufficient to decode the objects’ categories. Indeed, anterior regions of the parietal cortex are particularly sensitive to object elongation because it facilitates grasping (Chen et al., 2017). This possibility is also unlikely because all four categories contained elongated objects and our decoding results did not differ when tools, the category with the greatest number of elongated objects, was removed.

Another possibility is that that the dorsal pathway accomplishes categorization by computing an object’s structural description and then transmitting this information to the ventral pathway for recognition (Romei et al., 2011; Zachariou et al., 2017; Zaretskaya et al., 2013). Structural descriptions are a representation of ‘global shape’, which describe the spatial relations between an object’s parts, without describing the appearance of the parts themselves (Barenholtz & Tarr, 2006; Biederman, 1987; Hummel, 2000). Structural descriptions are crucial for basic-level categorization because exemplars within a category share the same structure, while varying in their individual features (Ayzenberg & Lourenco, 2022; Ayzenberg & Lourenco, 2019; Rosch et al., 2004; Tarr & Bülthoff, 1995). The dorsal pathway may be ideally suited to compute such a representation given its well-known role in computing spatial relations more generally. Indeed, recent studies have found that regions in PPC were functionally selective for an object’s structure, independent of other properties represented by the dorsal pathway, such as tools (Ayzenberg & Behrmann, 2022b). These PPC regions were capable of decoding an object’s category and mediated representations of structure in the ventral pathway (Ayzenberg et al., 2022). As mentioned previously, such structural information is readily transmitted via magnocellular input, providing a mechanism by which the dorsal pathway rapidly gains access to identity-relevant object information (Wang et al., 2022). Thus, not only might the dorsal pathway compute the spatial relations between objects to support action, but it may also compute the relations between object parts so as to form more complete representations of shape, in conjunction with ventral pathway (Ayzenberg & Behrmann, 2022a).

In summary, our results expand our understanding of the interactions between dorsal and ventral visual pathways. Although it is increasingly accepted that the two visual pathways interact, the precise nature of these interactions remain poorly understood. Here, we showed that identity-relevant object processing in the dorsal pathway precedes and predicts processing in the ventral pathway. Together, these findings provide an expanded understanding of the biological network that support visual object recognition.

## Acknowledgements

We are very grateful to Drs G. Gurariy and G. Caplovitz, and their co-authors, for providing us with the EEG data and the stimuli from their study, and their helpful comments. The current research was supported by a grant from the National Science Foundation to MB (BCS2123069) and from the National Institute of General Medical Sciences (NIGMS) to CS (T32GM081760). The content is solely the responsibility of the authors and does not necessarily represent the official views of the NIGMS, NSF, CMU or the University of Pittsburgh.

